# Clustering large-scale biomedical data to model dynamic accumulation processes in disease progression and anti-microbial resistance evolution

**DOI:** 10.1101/2024.09.19.613871

**Authors:** Kazeem A. Dauda, Olav N. L. Aga, Iain G. Johnston

## Abstract

Accumulation modelling uses machine learning to discover the dynamics by which systems acquire discrete features over time. Many systems of biomedical interest show such dynamics: from bacteria acquiring resistances to sets of drugs, to patients acquiring symptoms during the course of progressive disease. Existing approaches for accumulation modelling are typically limited either in the number of features they consider or their ability to characterise interactions between these features – a limitation for the large-scale genetic and/or phenotypic datasets often found in modern biomedical applications. Here, we demonstrate how clustering can make such large-scale datasets tractable for powerful accumulation modelling approaches. Clustering resolves issues of sparsity and high dimensionality in datasets but complicates the intepretation of the inferred dynamics, especially if observations are not independent. Focussing on hypercubic hidden Markov models (HyperHMM), we introduce several approaches for interpreting, estimating, and bounding the results of the dynamics in these cases and show how biomedical insight can be gained in such cases. We demonstrate this ‘Cluster-based HyperHMM’ (CHyperHMM) pipeline for synthetic data, clinical data on disease progression in severe malaria, and genomic data for anti-microbial resistance evolution in *Klebsiella pneumoniae*, reflecting two global health threats.

## Introduction

Accumulation models infer the dynamics by which features are acquired by a system over time [Diaz-Uriarte and Herrera-Nieto, 2022, Diaz-Colunga and Diaz-Uriarte, 2021, Diaz-Uriarte and Johnston, 2023]. A broad range of systems throughout biomedical sciences have been studied in this way. The most common is cancer progression, where genetic features (often mutations) are acquired as a tumour evolves. Accumulation modelling has also been applied to anti-microbial resistance, where features are either genes linked to drug resistance or drug resistance phenotypes [Aga et al., 2024, Greenbury et al., 2020, Moen and Johnston, 2022,Beerenwinkel et al., 2005], and to disease progression, where features are symptoms acquired by patients over time [Johnston et al., 2019, Schill et al., 2024]. In all these cases, accumulation modelling allows the dynamics of the underlying accumulation process to be inferred, predictions about the future behaviour of the system to be made, and mechanisms (for example, interactions between features) to be characterised using large-scale data [Aga et al., 2024, Diaz-Uriarte and Herrera-Nieto, 2022, Diaz-Uriarte and Johnston, 2023].

Particularly in the field of cancer progression, numerous approaches for accumulation modelling have been developed [Diaz-Uriarte and Johnston, 2023]. These include deterministic dependence models, developed to explore patterns of dependencies in the irreversible acquisition of binary traits using cross-sectional data [Diaz-Uriarte and Johnston, 2023]. These models include Oncogenetic trees (OT) [Desper et al., 1999, Szabo and Boucher, 2008], OncoBN (DBN) [Nicol et al., 2021], Conjunctive Bayesian Networks (CBN) [Gerstung et al., 2009,Montazeri et al., 2016], and Hidden Extended Suppes-Bayes Causal Networks (H-ESBCN, PMCE) [Angaroni et al., 2021], respectively. Several of these cancer progression models rely on the assumption that each subject provides a single data point and, as such, is independently observed. These represent crosssectional data, which is a standard format for the cancer progression models. However, these models do not incorporate essential evolutionary assumptions (e.g. species may share a common ancestor, heritable traits, etc.) and interpretations [Diaz-Uriarte and Johnston, 2023]. More recently, two changes to model structures have been introduced: stochastic rather than deterministic dependencies, where a feature can positively or negatively influence the acquisition probability of other features, realised in approaches like hypercubic transition path sampling (HyperTraPS) [Johnston and Williams, 2016, Greenbury et al., 2020], hypercubic Hidden Markov Models (HyperHMM) [Moen and Johnston, 2022] and mutual hazard networks (MHN) [Schill et al., 2019]; and support for phylogenetic relationships between observations, avoiding the issue of pseudoreplication [Maddison and FitzJohn, 2015] when related samples are assumed to be independent [Johnston and Williams, 2016, Greenbury et al., 2020, Moen and Johnston, 2022, Aga et al., 2024, Luo et al., 2023].

Such methods often impose limitations on how different features influence each other. For example, the *L*^2^ HyperTraPS model and equivalent mutual hazard networks allow every feature to influence every other’s acquisition probability, but no further influences (for example, pairs acting non-additively). HyperHMM (and the corresponding model structure of HyperTraPS-CT [Aga et al., 2024]) relaxes this restriction, allowing a feature’s acquisition rate to depend on arbitrary combinations of subsets of other features: every transition between two states has an independent transition probability. But it is limited in the number of features it can consider, with *>* 20 or so features being intractable. This reflects a general tradeoff in accumulation models: the size of the parameter space involved means that either the number of features involved, or the order of their interactions, is computationally limited.

Here, we propose a possible resolution to this tradeoff: dimensionality reduction through clustering features. In brief, we aim to group features whose behaviours are similar across the dataset, and work with a transformed dataset describing the presence or absence of members of feature sets, rather than individual features. This approach has the capacity to deal both with size and sparsity in large-scale datasets, although the transformation process introduces several protocol choices and corresponding complications in interpretation, which we will demonstrate and discuss. We place particular applied emphasis on two health challenges. The first is disease progression in severe malaria, a disease involving hundreds of millions of cases and hundreds of thousands of deaths every year, with a particular toll in the developing world [World Health Organization, 2023]. The heterogeneity of malaria progression poses clinical challenges, as optimal triage and treatment of patients with diverse profiles of symptoms may require understanding of the dynamics of the disease and their prognostic implications [Johnston et al., 2019]. The second is the acquisition of anti-microbial resistance (AMR)in *Klebsiella pneumoniae* (Kp), a pathogen of worldwide health concern responsible for many hospital-based infections [Effah et al., 2020, Navon-Venezia et al., 2017, Holt et al., 2015, Wyres et al., 2020]. The process of AMR evolution can be viewed as a stochastic acquisition of binary traits consisting of the presence or absence of genetic features [Greenbury et al., 2020, Baquero et al., 2021], and large-scale datasets allow this process to be explored in unprecedented detail [Aslam et al., 2021, Baker et al., 2023]. However, genomic data often characterises hundreds of genetic features of interest (individual mutations and/or complete genes acquired horizontally) for AMR, and an individual isolate will only have a small subset of these features. The joint issues of feature set size and sparsity feature prominently in the accumulation modelling of both malarial symptoms and genomic AMR data; we will proceed by establishing and exploring a collection of clustering methods as pre-processing steps before illustrating HyperHMM inference on the reduced-dimension datasets.

## Methodology

The idea of Cluster-based HyperHMM (CHyperHMM) is to cluster datasets with large numbers of features *p*, to reduce dimensionality to a smaller set of *p*_*C*_ ‘effective features’ or ‘feature clusters’. HyperHMM, or other accumulation modelling approaches, can then be used to learn the dynamics by which the system acquires elements of these effective features (Fig. 1). But in amalgamating features into these clusters, several points must be considered. First, different clustering methods can be used to assign clusters based on which features appear in similar patterns across observations. Second, once clusters are assigned (i.e. which features correspond to which cluster), an occupancy protocol must be used. For example, if an observation has some but not all features of cluster *X*, does this count as having acquired cluster *X* or not? Third, the relationship, if any, between observations must be considered. If they are fully independent, this is not an issue. But if they are, for example, linked by a shared ancestry, the possibility of similarity-by-descent should be addressed to avoid ‘pseudoreplication’ – assigning undue weight to observations that are not independent. Fourth, we must consider how to interpret the outputs of accumulation modelling based on feature clusters rather than individual features, particularly given that different choices may be made for the previous cases. **Clustering methods**. We cluster features based on their appearance across samples in the dataset – hence the task is to cluster binary strings by similarity. A wider range of clustering choices exists in the literature [Charrad et al., 2014, Ren et al., 2021, Botelho et al., 2023, Batool and Hennig, 2021, Patel et al., 2022, Ezugwu et al., 2022, Abualigah et al., 2018]. For simplicity, reproducibility, and interpretability, we use either k-means clustering ([Ikotun and Ezugwu, 2022,Ding and He, 2004,MacQueen, 1967]) using Manhattan distance and the gap-statistic method for determining the optimal number of clusters ([Tibshirani et al., 2002]), or the validity index (based on scattering within, and separation between, clusters) [Halkidi et al., 2000]; other methods are of course possible ([Charrad et al., 2014], see Software Implementation). Clusters are visualised using principal components analysis [Lever and Altman, 2017].

**Figure 1.**
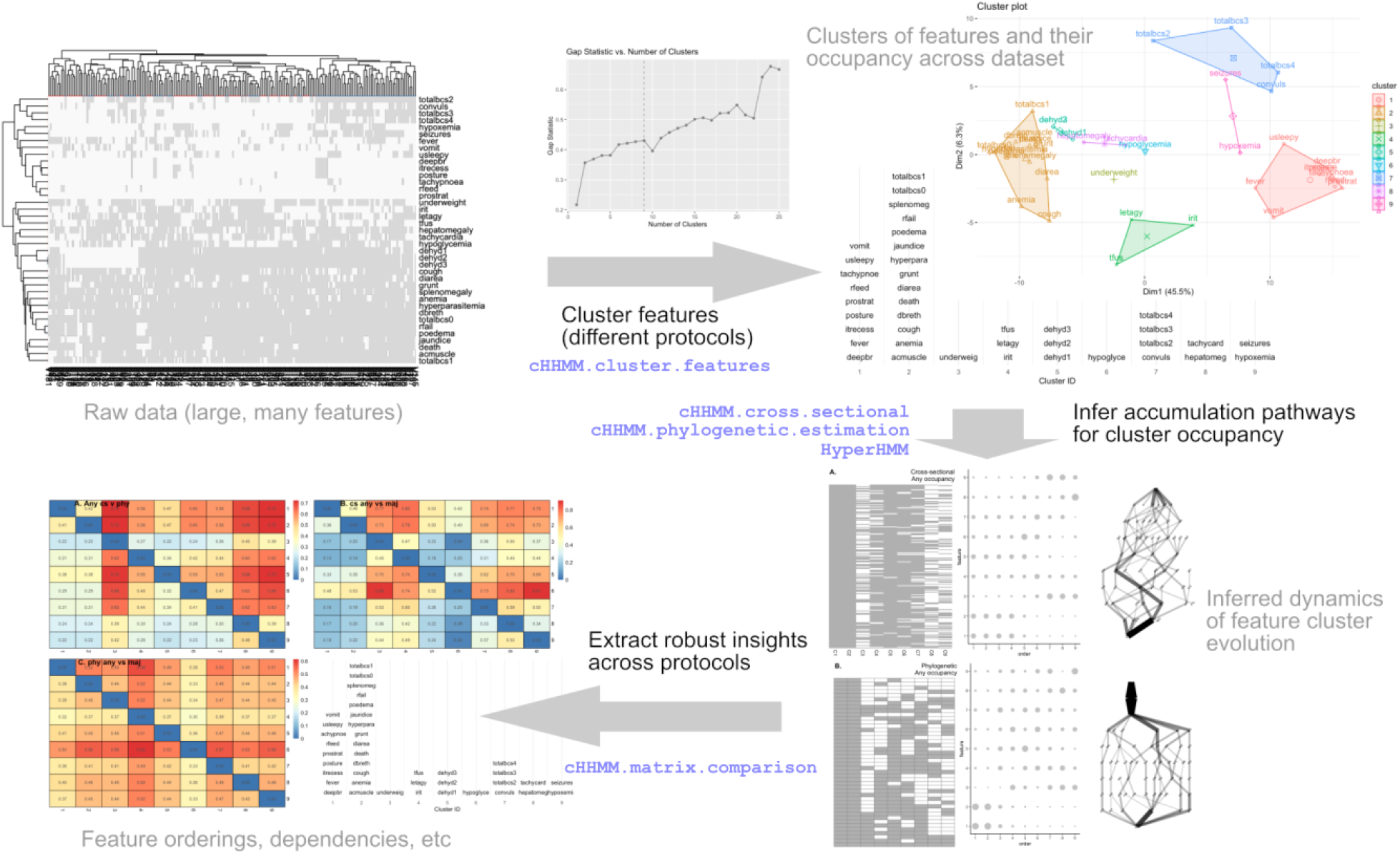
Overview of CHyperHMM inference. The raw data consists of a large number of features (presence (1) and absence (0) of AMR genes, referred to as the ‘barcode’). We then cluster the features using different occupancy protocols to generate a data structure acceptable for the HyperHMM method. To infer the cumulative pathways of the feature clusters, independently or phylogenetically, and across different protocols, we use HyperHMM and evaluate the robustness of these protocols using cHMM matrix comparison.

### Cluster occupancy assignment

Once the member features of each cluster are determined, the next task is to assign a presence or absence marker to each cluster for each sample. In general, samples may possess a subset of features for each cluster, and a protocol is required for determining which feature subsets correspond to cluster presence. While the most interpretable protocol will depend on scientific application, we consider a spectrum of options here: (i) at least one member feature; (ii) *≥* 50% of member features; (iii) *> c* member features, where *c* is the mean number of member features present across samples.

### Relationships between observations

In some cases, the relationship between samples in the dataset may be well known. For example, they may be genuinely independent (symptom profiles from different patients), or a phylogeny relating the observations may be independently known. However, if observations are known to be related but the relationship is unknown, we must consider how to proceed. Here, we consider three cases which in a sense reflect the limiting bounds of relatedness. First, we assume complete independence, which will lead to pseudoreplication if the observations are in fact related: the effective sample size will be lower than the number of observations, because several arise from a more limited number of common ancestors. Second, we construct an estimated simple phylogeny linking the observations, by clustering the dataset and extracting the dendrogram reflecting the cluster structure. This attempts to capture similarity by descent, simply by relating the most similar observations throughout the dataset. Third, we use a maximum parsimony (MP) approach from evolutionary biology to estimate and describe the phylogenetic relatedness of the observations. The MP method is robust in identifying tree structures that require the minimum number of evolutionary changes to account for the observations as the tips of the tree.

### Accumulation modelling with HyperHMM

Given a particular set of choices for the protocols above, a given dataset will be represented as a set of *n* observations of *p*_*C*_ binary markers corresponding to cluster presence/absence, and a phylogeny linking these observations (which may be the trivial ‘star’ phylogeny reflecting complete independence). This data structure is the input to accumulation modelling approaches like HyperHMM. That algorithm proceeds as follows [Moen and Johnston, 2022]. First, ancestral states on the phylogeny are reconstructed using a simple logic rule reflecting parsimonious accumulation (in other words, irreversible) dynamics: if two daughers share a presence marker for a given feature, their ancestor is assigned a presence marker, otherwise the ancestor is assigned an absence marker. Next, transitions between states from ancestor to descendant are collected from the labelled phylogeny. These before-after transitions are the raw data for HyperHMM. The algorithm itself is an adapted Baum-Welch (expectation maximisation) algorithm estimating maximum-likelihood weights for a hypercubic transition graph reflecting all possible transitions between states.

### Comparing outputs

Accumulation models fitted using different choices of the protocols above will in general give different output predictions. Methods for comparing accumulation model outputs are established for tree-based outputs, but less so for transition networks resulting from stochastic models [Pascual et al., 2024]. We will use one approach focussing on [*A*_*ij*_], the matrix of probabilities that feature *i* is acquired in ordinal timestep *j*. All accumulation models will explicitly or implicitly output this information. We can 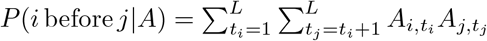 compute *P* (*i* before *j*|*A*) = ^*L L*^ *A*_*i,t*_ *A*_*j,t*_. We can then compare and/or combine *P* (*i* before *j*|*A*) with *P* (*i* before *j*|*B*) (and others) in various ways; an example is to compute

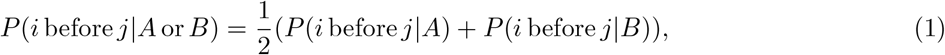

giving the overall probability that feature *i* is acquired before feature *j* while weighting the two outputs *A* and *B* equally. When we are uncertain of the best protocol to use (for example, for occupancy rules), this approach will report the general pairwise ordering dynamics that are more or less robustly observed across different protocol choices.

### Malaria clinical data

Clinical symptom data from Gambian children with severe malaria (SM) was used from [Johnston et al., 2019]. Quoting that article: ‘The study population consisted of 2915 children aged 4 months to 15 years diagnosed with SM according to the WHO definition. Children were admitted to the Royal Victoria Teaching Hospital (RVTH), Banjul, The Gambia from January 1997 to December 2009. The study was originally designed to study genetic variants associated with SM. The initial set of variables used for feature selection included those present in the case report form… Children aged 4 months to 15 years were eligible for enrolment if they had a blood smear positive for asexual *Plasmodium falciparum* parasites and met one or more WHO criteria for SM: Coma (assessed by the British Coma Score, BCS), severe anaemia (haemoglobin [Hb]*<*50g/L or packed cell volume [PCV]*<*15), RD (costal indrawing, use of accessory muscles, nasal flaring, deep breathing), hypoglycaemia (2.2mM), decompensated shock (systolic blood pressure less than 70mmHg), repeated convulsions (*>*3 during a 24-h period), acidosis (plasma bicarbonate *<*15mmol/L) and hyperlactatemia (plasma lactate *>*5mmol/L). CM was defined as a BCS of 2 or less with any *P. falciparum* parasite density. SMA was defined as haemoglobin under 50g/L. Hepatomegaly was defined as *>* 2cm of palpable liver below the right costal margin. Patients were enrolled in the study if written informed consent was given by the parent or guardian. The study protocol was approved by the Joint Gambia Government/MRC Ethical Committee (protocol numbers 630 and 670).

***Klebsiella pneumoniae* genomic data**. A dataset containing Kleborate output and metadata for 47721 whole-genome sequenced Kp isolates was extracted and curated from publically available data [Lam et al., 2021, Sánchez-Busó et al., 2022, Argimón et al., 2021]. We downloaded two datasets from pathogen-watch through https://pathogen.watch. One dataset contained metadata such as the continent, country, and host organism; the other contained with Kleborate output for all samples. A total of 17 columns containing acquired antibiotic resistance genes (ARGs) were selected from the Kleborate output. Kleborate groups these genes by predicted phenotypic resistance, and we further transformed them into wide format with 481 genotypic features. The features were binarized to indicate the presence (1) or absence (0) of a specific gene, which is required by our developed method. The cleaning steps removed a collection of symbols and patterns describing fine-grained aspects of the data, such as ‘+13V’ (description of specific variant), asterisks, carets, and question marks (describing the extent of incomplete nucleotide matching between an identified genetic feature and the corresponding reference sequence).

### Software implementation

We use R [R Core Team, 2022] with libraries: `ggplot2` [Wickham, 2016], `ggpubr` [Kassambara, 2023], `pheatmap` [Kolde, 2019], `ggplotify` [Yu, 2023], `igraph` [Csardi and Nepusz, 2006], `ggraph` [Pedersen, 2021], and latticeExtra [Sarkar and Andrews, 2022] for visualisation; `cluster` [Maechler et al., 2022], `factoextra` [Kassambara and Mundt, 2020], `NbClust` [Charrad et al., 2014] for clustering and multivariate analysis; `phytools` [Revell, 2024], `phangorn` [Schliep, 2011, Schliep et al., 2017], `ape` [Paradis and Schliep, 2019] for processing and manipulating phylogenies; and `parameters` [Lüdecke et al., 2020], `R.utils` [Bengtsson, 2022], `stringr` [Wickham, 2022], and `git2r` [git2r Authors, 2023] for the pipeline and workflow. Of particular note, the `NbClust` package implements a wide variety of methods for clustering and choice of cluster number [Charrad et al., 2014]. We also use HyperHMM code from https://github.com/StochasticBiology/hypercube-hmm [Moen and Johnston, 2022]. CHyperHMM code is freely available at https://github.com/Dydx1989/Cluster-HyperHMM.

## Results

### Identifying accumulation pathways from synthetic data

We first use a simple synthetic dataset with *n* = 200, *p* = 160 to demonstrate CHyperHMM. The data are constructed from a generating mechanism of two mutually exclusive evolutionary pathways, testing the ability of the approach to resolve negative interactions between features. There are four sets of ‘super-features’, each of which corresponds to a stochastic ‘up-regulation’ of a set of features; another set of features is sparsely occupied at random with no link to the super-features (Fig. 2A). The super-features evolve down one of the mutually exclusive pathways 0000-0001-0011-0111-1111 or 0000-1000-1100-1110-1111. Fig. 2B-C demonstrate the clustering of these data with the gap statistic behaviour, identifying *p*_*C*_ = 4 clusters corresponding to the governing ‘super-features’. Fig. 2D shows HyperHMM inference of the accumulation dynamics of these clusters, capturing the mutually exclusive pathways governing the system’s behaviour (given the arbitrary cluster labelling, these are 2-3-4-1 and 1-4-3-2).

**Figure 2.**
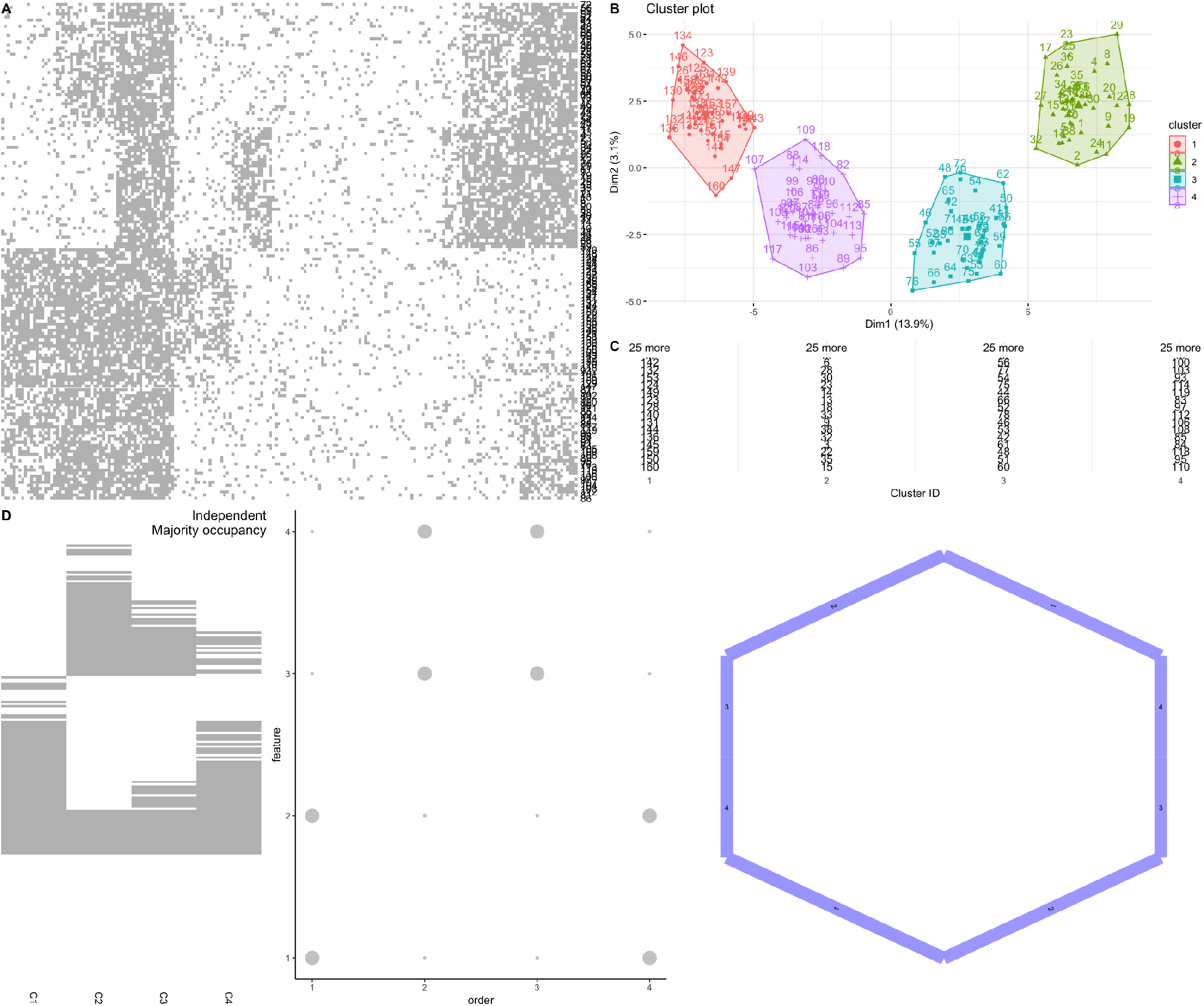
Synthetic data case study. (A) Raw (simulated) data with *n* = 200, *p* = 160; observations are columns, with presence (grey) and absence (white) for each feature (row). (B-C) Clustering features with k-means and gap statistic. (B) Principal components analysis embedding of feature profiles across observations, with clusters illustrated by convex hull; (C) partial membership lists of the features in each cluster. (D) HyperHMM inference on the clustered dataset (left), assuming independent observations and using the ‘majority’ occupancy rule (a cluster is ‘present’ if more than half its constituent features are present). (centre) Ordering plot giving the probability (point size) that a feature (row) is acquired at a given ordering (column) in the evolutionary process. (right) Inferred transition graph, with the all-absent state in the top centre and transitions proceeding down the graph. Edge weights give probabilities; edge labels give the acquired feature (cluster) for each transition.

### Severe malaria disease progression

We next considered a dataset on disease progression, specifically the symptoms exhibited by children with severe malaria ([Johnston et al., 2019]; see Methods). After removing outcome and intervention markers (death and transfusion) and interpreting missing data as absence markers, *p* = 37 symptoms remained for each of *n* = 2914 individuals (Fig. 3A). Through the ‘nbclust’ approach, the malaria symptom set is assigned *p*_*C*_ = 13 clusters (Fig. 3B-C). Cluster 7 has the most member features, including many rare symptoms in the dataset: severe coma, respiratory failure, altered posture, and several more. Cluster 13 has several, more common member features, including cough, anemia, diarrhea, and low weight. Several other clusters (1-4, 6, 8, 10) have only a single member feature, occupying intermediate positions in the first principal component of cross-patient variation but distinct enough to be assigned their own identities. Other clusters group symptoms intuitively: for example, cluster 11 contains the less severe coma markers, and cluster 9 contains deep breathing and breathing using accessory muscles.

**Figure 3.**
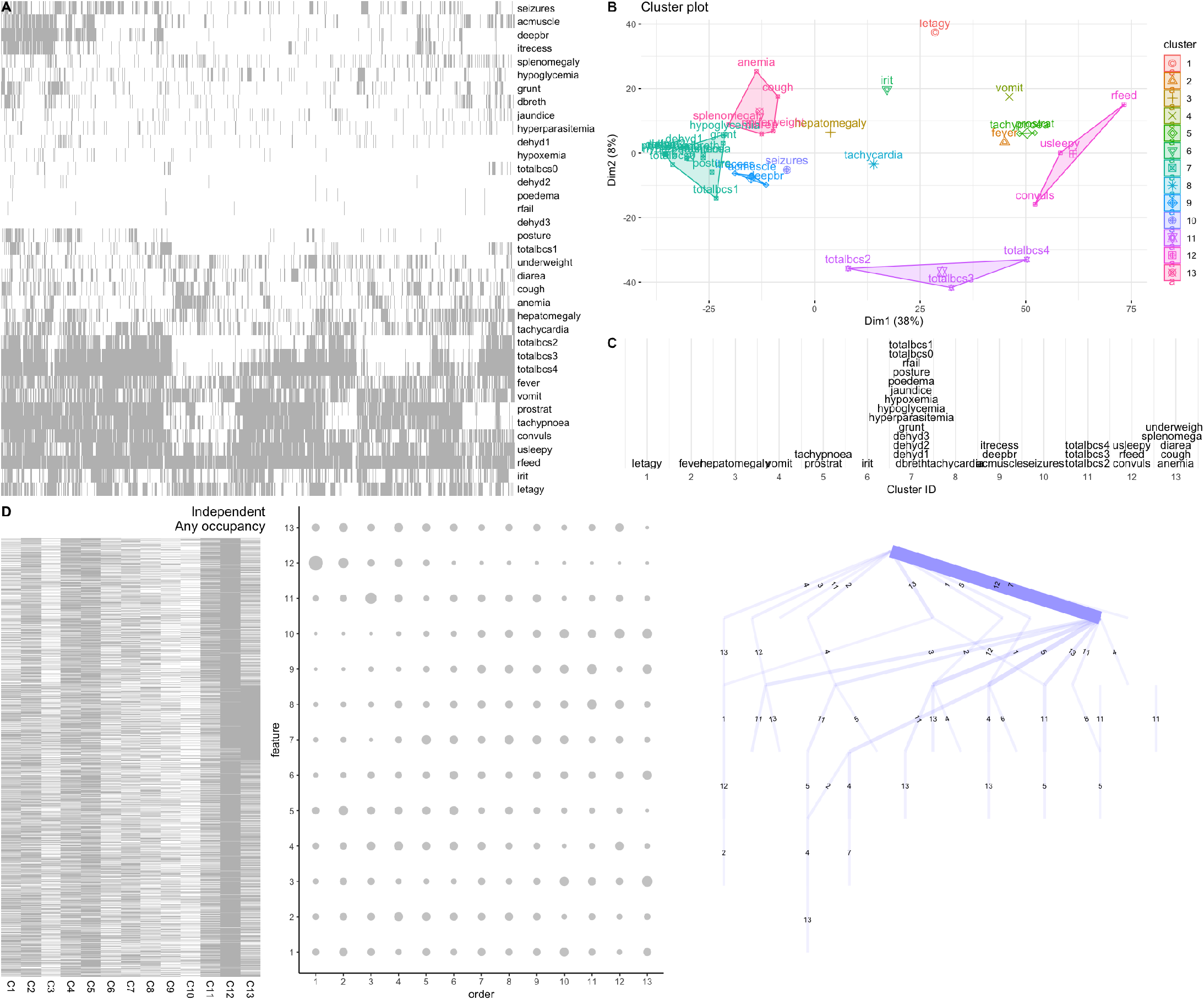
Severe malaria case study. Structure as in Fig. 2. (A) Patient data with presence (grey) and absence (white) for each symptom (row); symptom codes are given in Supp. Table S2. (B) Cluster structure in PCA space and (C) membership of each cluster from k-means clustering with the validity index for selecting cluster number. (D) HyperHMM inference on the clustered dataset (left), assuming independent observations and using the ‘any’ occupancy rule (a cluster is ‘present’ if any of its constituent features are present). (centre) Ordering plot giving the probability (point size) that a feature (row) is acquired at a given ordering (column) in the evolutionary process. (right) Inferred transition graph, with the all-absent state in the top centre and transitions proceeding down the graph. Edge weights give probabilities; edge labels give the acquired feature (cluster) for each transition.

Due to the independent nature of disease progression in individual patients (all of whom start independently, symptom-free), we use the cross-sectional realisation of these data, and choose the ‘any’ occupancy rule due to the common mechanistic relatedness of symptoms within a cluster (alternative choices are shown in Supp. Fig. S1.). With this choice (Fig. 3D), occupancy of cluster 12 (unusual sleepiness, reduced feeding, and convulsions) is almost complete, and it is inferred to be the likely first acquired cluster. Cluster 13 (several common features), cluster 1 (lethargy) and cluster 5 (tachypnoea and prostration) are likely next acquired features. Cluster 11 (coma score between 2 and 4) is the likely third step, with clusters 4 and 2 (vomiting and fever) likely appearing next. Patterns involving all these acquisitions and more are less common, so the pathways of disease progression following this set are less canalised, but clusters 3 (hepatomegaly), 6 (irritability), 9 (three breathing-related symptoms), and 10 (seizures) are inferred to be acquired later.

If the ‘majority’ occupancy rule is enforced (Fig. 3), inferred orderings and pathways do not differ hugely – perhaps unsurprisingly, given that for the several single-feature clusters, ‘any’ and ‘majority’ occupancy rules are identical. Cluster 7, the larger collection of relatively rare symptoms, has a higher probability of late acquisition under the majority rule. This is because while a patient may exhibit a small number of rare symptoms, patients exhibiting more than half of this set is almost never observed. The comparison matrix approach (see Methods; Supp. Fig. S2) emphasises that early acquisition of cluster 12 (and cluster 5) are robust across occupancy protocol choices; late acquisitions of 7, 9, 10 are also generally supported regardless of protocol. Although not of central interest for this case study (as the individual patient samples are known to be independent), Supp. Fig. S2 also shows that these observations are largely robust if the observations are assumed to be related by similarity rather than truly independent.

Inference using the full feature set using HyperTraPS [Johnston et al., 2019] – taking over an order of magnitude more time – is consistent with many of these observations. The symptoms in cluster 12 are among the earliest inferred, and cluster 5 follow soon thereafter. Fever, vomiting, lethargy, and limited coma scores are all inferred early symptoms; hepatomagaly, seizure, irritability are inferred as intermediate acquisitions; the rarer members of cluster 7 appear together as inferred late symptoms.

### Acquisition of AMR genetic features in *Klebsiella pneumoniae*

We next explored a dataset from Pathogenwatch on genetic features associated with AMR in Kp [Sánchez-Busó et al., 2022, Argimón et al., 2021] (see Methods). The distribution of the *n* = 41089 sequenced isolates across continents is as follows: Africa 2327 (4.88%); Americas (North and South America) 13300 (27.87%); Asia 12200 (25.57%); Europe 17941 (37.60%); and Oceania 1953 (4.09%).

The dataset involves *p* = 481 genetic features, including chromosomal and plasmid gene acquisitions. Fig. 4A immediately demonstrates some of the features of this dataset. It is very sparse, with most features present in only a small number of isolates – but also heterogeneous, with some much more common features. Several immediate advantages of the clustering approach, suggesting *p*_*C*_ = 8 clusters, are clear from its behaviour (Fig. 4B). The many rare features that contribute to the data sparsity are clustered together into a very large single feature (akin to the many rare symptoms in the malaria case study). These rare features, while certainly individually interesting, can then in a sense be ‘factored out’ of the analysis of the remaining features which exhibit more structured dynamics. The presence of this large feature (cluster 4, in the arbitrary labelling) does not interfere with assignment of rare features that nonetheless exhibit structure: cluster 5 overlaps with cluster 4 but contains many fewer features which internally covary. The first principal component of the feature embedding is tightly linked to sparsity: the clusters most distal from cluster 4 involve features like *CTX-M-15, TEM-1D, sul2, strA, strB* which are among the most common in the dataset. The clustering approach identifies subsets of these more represented features with similar profiles.

**Figure 4.**
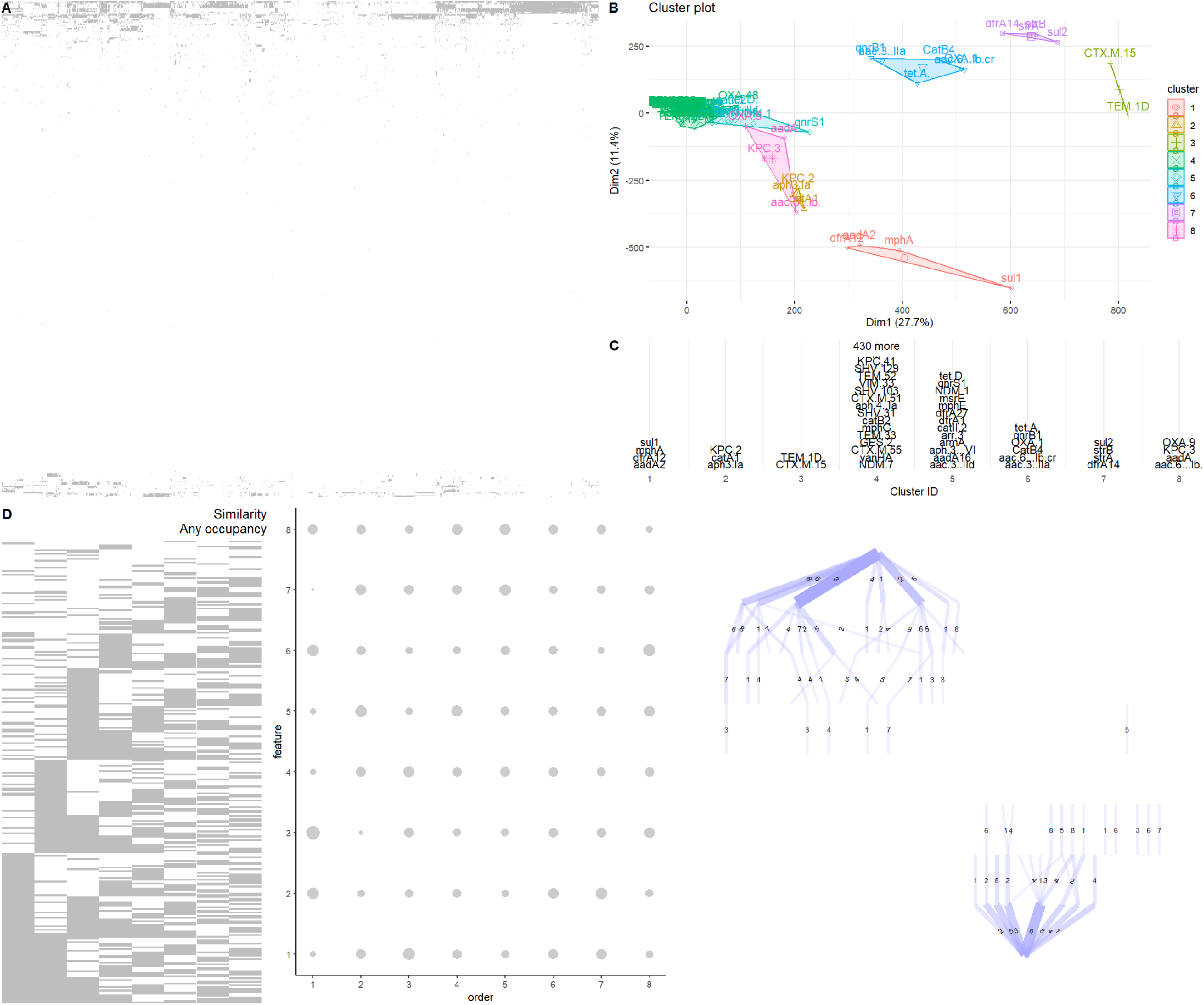
Klebsiella AMR case study. Structure as in Fig. 2. (A) *Klebsiella pneumoniae* isolate data with presence (grey) and absence (white) for each mutation (rows; names omitted for clarity). The dataset is very sparse, with most mutations very rarely present, but with some mutations much more common (highest and lowest rows) and correlated. (B) Cluster structure in PCA space and (C) membership of each cluster from k-means clustering with the gap statistic for selecting cluster number. Non-alphanumeric characters in feature names are replaced by periods. (D) HyperHMM inference on the clustered dataset (left), using a similarity measure to estimate the phylogenetic relationship between observations and and using the ‘any’ occupancy rule (a cluster is ‘present’ if any of its constituent features are present). (centre) Ordering plot giving the probability (point size) that a feature (row) is acquired at a given ordering (column) in the evolutionary process. (right) Inferred transition graph, with the all-absent state in the top centre and transitions proceeding down the graph. Edge weights give probabilities; edge labels give the acquired feature (cluster) for each transition.

We examined the function and provenance of member genes of all clusters except the giant sparse cluster 4 using the Comprehensive Antibiotic Resistance Database (CARD) [Alcock et al., 2023]. The overall trend in this dataset is antibiotic inactivation as the mode (Supp. Table S2), with features being more prevalent in plasmid form over chromosomal, but present in both across all samples (Supp. Fig. S3). The genes within most clusters are mixed in terms of both mechanism and antibiotic classes they confer resistance to (Supp. Table S2). The most common mechanism in the clusters are inactivating the antibiotic agent. Three clusters (2,3 and 8) consists solely of genes that inactivate the agent; the two genes in cluster 3 are the most consistent with respect to antibiotic class.

We begin the inference process by using a similarity picture to estimate a phylogenetic relationship linking the isolates, and consider the ‘any’ occupancy rule. The output of inference then reports the evolutionary pathways by which isolates acquire at least one mutation from each cluster, assuming that more similar AMR profiles are more related. The most likely first step under this protocol is cluster 3 (*TEM-1D* or *CTX-M-15*), with reasonable probabilities also for cluster 6 (*aac3, aac6, catB, OXA-1, qnrB*, or *tetA*), cluster 2 (*aph(3’)-Ia, catA, KPC-2*), and cluster 8 (*aac6, aadA, KPC-3, OXA-9*). Clusters 1 (*aadA2, dfrA12, mphA, sul1*), 7 (*dfrA14, strA, strB, sul2*), and 5 (a larger collection) are likely next steps. Cluster 4, involving a set of over 430 relatively rare mutations, has a relatively uniform ordering distribution, suggesting that acquisitions within this range of rare feature can occur somewhat independently of the other mutations in the dataset.

To demonstrate the influence of different modelling assumptions, we also analysed these data using the ‘majority’ occupancy rule and assuming independence of isolates (Supp. Fig. S4). Using this protocol, as with the malaria case study, substantially shifts the acquisition ordering of the cluster of many rare features: cluster 4 is now very likely acquired last. Acquisition of cluster 3 (first in the ‘any’ case) is still early, but now is likely followed more immediately by clusters 1 and 7, with the 2/6/8 group that was acquired early in the ‘any’ case now appearing relatively later. A relatively clear modal path (3-7-6-1-5-8-2-4) in fact appears, but must be interpreted in light of the substantial capacity for pseudoreplication that this protocol allows. The consistent early appearance of cluster 3 across occupancy rules is reflected in the comparison matrix plots (Supp. Fig. S5). These plots also show that the proportion of pairwise comparisons which are robust to the choice of cross-sectional versus related observations is relatively low, emphasising the importance of accounting for relatedness in datasets like this.

Non-trivial structures are observed throughout the ordering plots in the Klebsiella case study. Bimodal and multimodal ordering distributions – where a feature has a high early acquisition probability and a high late acquisition probability, but lower probability at intermediate times – are characteristic of separate evolutionary pathways. This bimodality is observed, for example, with cluster 5 in the ‘majority’ case and to some extent for cluster 3 in the ‘any’ case. This latter case involves a strikingly low probability that cluster 3 is the *second* feature to be acquired – supporting the above picture that it may share some influential relationship with other clusters.

We also asked how these inferred dynamics might differ between different regions. To this end, we performed inference separately for subsets of data from different continents (Fig. S6). Here, the broader aspects of the dynamics mentioned above – early acquisition of cluster 3 and 6, with some multimodality – are observed at the continent level too. Notably, in the Americas and Europe, cluster 3 is less likely as a first acquisition, instead appearing more commonly after an alternative, and cluster 1 is somewhat more likely to appear early. As cluster 3 involves two betalactamase elements acquired via plasmid transfer, this could indicate that acquisition of plasmids containing these specific elements is less common in these continents. However, the ordering distributions at a continent scale typically demonstrate a spread of probability for each cluster, so these effects certainly do not constitute universal rules.

## Discussion

The family of approaches for addressing ‘accumulation modelling’ problems is increasing over time, with developments in the cancer progression field, evolutionary biology, and more diverse applications [Diaz-Uriarte and Johnston, 2023]. A tension that exists across these applications is between the number of features that can be modelling and the flexibility of interactions that can be inferred between them. The HyperHMM picture supports stochastic positive and negative dependencies between not just individual features but arbitrary subsets of features [Moen and Johnston, 2022]; as every edge on the hypercubic transition graph is indepedently parameterised, all subsets of features can influence the acquisition of every other. This is arguably the most flexible picture possible without involving reversibility (which is captured in, for example, [Johnston and Diaz-Uriarte, 2024], but severely limits feature set size).

This flexibility constrains HyperHMM and comparable approaches to limited numbers of features. Here we have suggested that clustering to reduce large feature sets to smaller sets of feature collections can provide a route of enquiry in the face of these constraints. In the case of sparse datasets, clustering arguably focuses on the more interesting feature sets – those more represented in the data (potentially increasing the statistical power available for their accurate description) and which covary in non-trivial ways. In each of our case studies we saw that features are clustered intuitively (for example, malaria symptoms of coma severity cluster together, as do symptoms involving breathing phenotypes). Certainly information is lost during this dimensionality reduction, but general trends in the accumulation dynamics – as shown by the synthetic and malaria studies – can be identified at dramatically reduced computational cost.

This advantage comes with the disadvantage that the outputs of inference are harder to interpret. We have attempted to demonstrate several different protocols to suit different scientific circumstances, and ways to seek results that are robust with respect to such protocol choice. However, it must of course be noted that there is a large number of methods both for unsupervised clustering of data and for choosing a number of clusters given a dataset [Xu and Wunsch, 2005, Saxena et al., 2017, Charrad et al., 2014], and different scientific circumstances may support the choice of different approaches to those we have presented here.

We have focussed on HyperHMM due to its flexible model structure, but of course there is no reason that clustering cannot be applied as a pre-processing step in other accumulation modelling approaches. The output, in the case of cross-sectional observations, fundamentally remains a collection of binary vector observations. HyperTraPS has recently been expanded to support the full subset-based influence structure of HyperHMM [Aga et al., 2024] but also supports simpler model structures (similar to MHN [Schill et al., 2019]); and other approaches from the cancer literature could also readily be used to analyse such data [Diaz-Uriarte and Johnston, 2023, Diaz-Uriarte and Herrera-Nieto, 2022]. In the case of observations linked by a phylogeny (known or estimated), HyperTraPS or TreeMHN [Luo et al., 2023] would be possible downstream alternatives.

## Acknowledgements

This work was supported by the Trond Mohn Foundation [project HyperEvol under grant agreement No. TMS2021TMT09], through the Centre for Antimicrobial Resistance in Western Norway (CAMRIA) [TMS2020TMT11]. This project has received funding from the European Research Council (ERC) under the European Union’s Horizon 2020 research and innovation programme [grant agreement No. 805046 (EvoConBiO)].

## Supplementary Material

**Figure S1.**
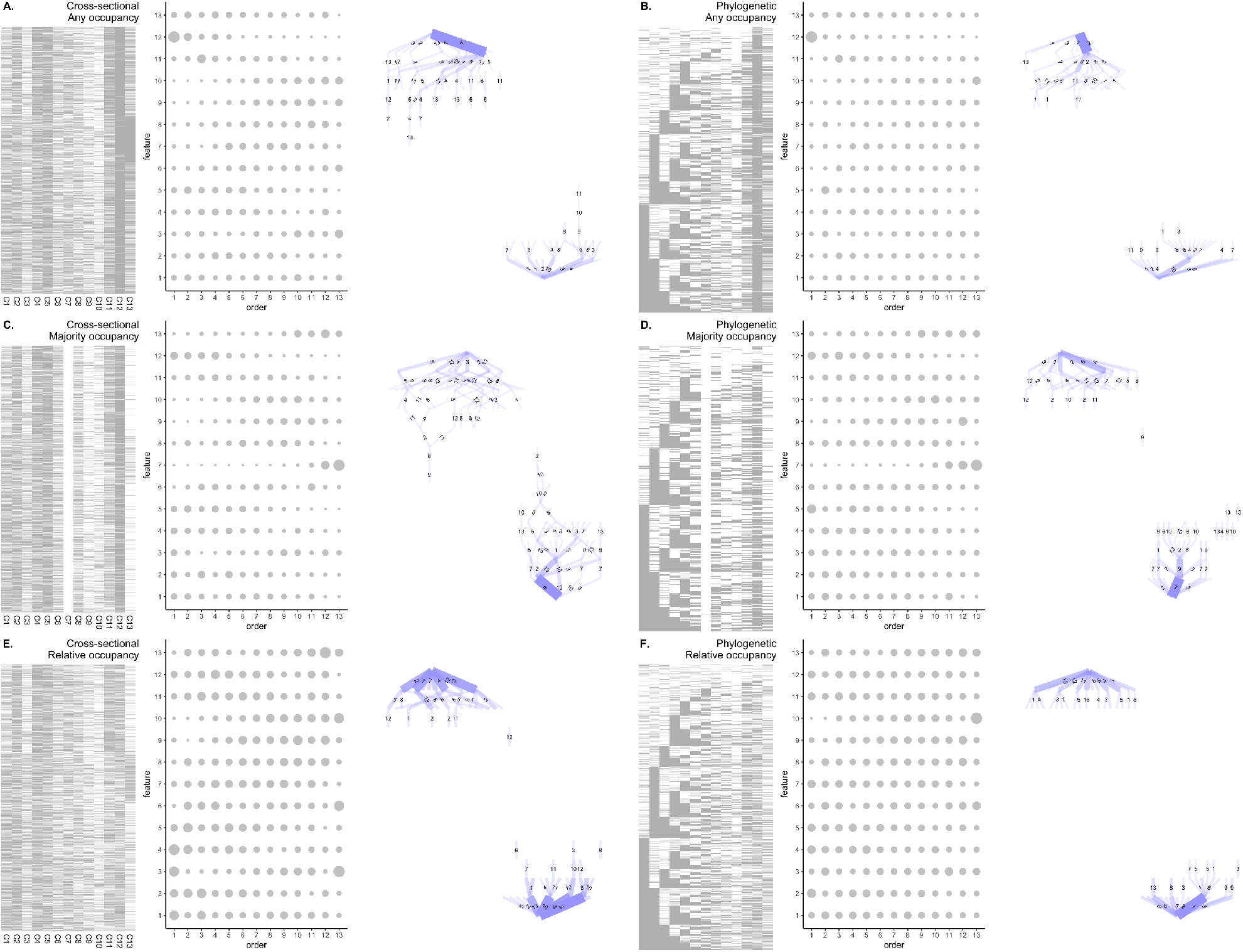
Severe malaria case study – alternative approaches. Post-clustering data, ordering matrices, and transition graphs (as in Main Text figures) for different protocol choices in the severe malaria dataset.

**Table S2.** Malaria symptom codes. Following [Johnston et al., 2019], a collection of binary-coded variables describing the presence or absence of different malaria features. Several are originally derived from continuous or ordinal statistics.

| Code | Symptom |
| --- | --- |
| cough | Coughing |
| deepbr | Deep breathing |
| convuls | Reported convulsions prior to admission |
| diarea | Diarrhea |
| vomit | Vomiting |
| usleepy | Unusually sleepy |
| irit | Irritable or restless |
| dbreth | Difficulty breathing |
| letagy | Lethargy |
| rfeed | Reduced feeding |
| jaundice | Jaundice |
| grunt | Grunting |
| acmuscle | Accessory muscles used in respiration |
| rfail | Renal failure |
| poedema | Pulmonary oedema |
| itrecess | Intercostal recession |
| fever | Fever |
| hypoglycemia | Hypoglycemia |
| hypoxemia | Hypoxemia |
| anemia | Anemia |
| hepatomegaly | Hepatomegaly |
| seizures | Seizures |
| hyperparasitemia | Hyperparasitemia |
| posture | Abnormal posturing |
| prostrat | Prostration |
| tachycardia | Tachycardia |
| tachypnoea | Tachypnoea |
| underweight | Underweight |
| splenomegaly | Splenomegaly |
| dehyd[X] | Dehydration score over $X$ |
| totalbcs[X] | Coma score under $X$ |

**Table S2.** Properties of genes in the non-giant Klebsiella cluster. Data extracted from CARD [Alcock et al., 2023]. Genes are listed by their primary name in CARD. *catB3, APH(6)-Ia* and *APH(6)-Id* are synonyms for *catB4, strA* and *strB* respectively. Genes *aph3-Ia* and *aph(3’)-Ia* were considered equivalent. For genes *catII, aph(3’)-VI* and *aac(6’)-Ib-cr* a member of those gene groups were looked up in CARD and presented in the table.

| Cluster | ARG | Gene family | Resistance mechanism | Antibiotic class |
| --- | --- | --- | --- | --- |
| 1 | sul1 | sulfonamide resistant sul | antibiotic target replacement | sulfonamide antibiotic |
| 1 | dfrA12 | trimethoprim resistant dihydrofolate reductase dfr | antibiotic target replacement | diaminopyrimidine antibiotic |
| 1 | aadA2 | ANT(3'') | antibiotic inactivation | aminoglycoside antibiotic |
| 1 | mphA | macrolide phosphotransferase (MPH) | antibiotic inactivation | macrolide antibiotic |
| 2 | APH(3')-Ia | APH(3') | antibiotic inactivation | aminoglycoside antibiotic |
| 2 | KPC-2 | KPC beta-lactamase | antibiotic inactivation | monobactam, carbapenem, cephalosporin, penam |
| 2 | catA1 | chloramphenicol acetyltransferase (CAT) | antibiotic inactivation | phenicol antibiotic |
| 3 | CTX-M-15 | CTX-M beta-lactamase | antibiotic inactivation | cephalosporin, penam |
| 3 | TEM-1 | TEM beta-lactamase | antibiotic inactivation | monobactam, cephalosporin, penam, penem |
| 5 | msrE | msr-type ABC-F protein | antibiotic target protection | macrolide antibiotic, streptogramin antibiotic |
| 5 | QnrS1 | quinolone resistance protein (qnr) | antibiotic target protection | fluoroquinolone antibiotic |
| 5 | arr-3 | rifampin ADP-ribosyltransferase (Arr) | antibiotic inactivation | rifamycin antibiotic |
| 5 | NDM-1 | NDM beta-lactamase | antibiotic inactivation | carbapenem, cephalosporin, cephamycin, penam |
| 5 | APH(3')-VIa | APH(3') | antibiotic inactivation | aminoglycoside antibiotic |
| 5 | tet(D) | major facilitator superfamily (MFS) antibiotic efflux pump | antibiotic efflux | tetracycline antibiotic |
| 5 | catII | chloramphenicol acetyltransferase (CAT) | antibiotic inactivation | phenicol antibiotic |
| 5 | armA | 16S rRNA methyltransferase (G1405) | antibiotic target alteration | aminoglycoside antibiotic |
| 5 | aadA16 | ANT(3'') | antibiotic inactivation | aminoglycoside antibiotic |
| 5 | dfrA1 | trimethoprim resistant dihydrofolate reductase dfr | antibiotic target replacement | diaminopyrimidine antibiotic |
| 5 | AAC(3)-IId | AAC(3) | antibiotic inactivation | aminoglycoside antibiotic |
| 5 | mphE | macrolide phosphotransferase (MPH) | antibiotic inactivation | macrolide antibiotic |
| 5 | dfrA27 | trimethoprim resistant dihydrofolate reductase dfr | antibiotic target replacement | diaminopyrimidine antibiotic |
| 6 | catB3 | chloramphenicol acetyltransferase (CAT) | antibiotic inactivation | phenicol antibiotic |
| 6 | AAC(3)-IIa | AAC(3) | antibiotic inactivation | aminoglycoside antibiotic |
| 6 | QnrB1 | quinolone resistance protein (qnr) | antibiotic target protection | fluoroquinolone antibiotic |
| 6 | AAC(6')-Ib-cr1 | AAC(6') | antibiotic inactivation | fluoroquinolone antibiotic, aminoglycoside antibiotic |
| 6 | tet(A) | major facilitator superfamily (MFS) antibiotic efflux pump | antibiotic efflux | tetracycline antibiotic |
| 6 | OXA-1 | OXA beta-lactamase | antibiotic inactivation | cephalosporin, penam |
| 7 | sul2 | sulfonamide resistant sul | antibiotic target replacement | sulfonamide antibiotic |
| 7 | APH(6)-Id | APH(6) | antibiotic inactivation | aminoglycoside antibiotic |
| 7 | APH(6)-Ia | APH(6) | antibiotic inactivation | aminoglycoside antibiotic |
| 7 | dfrA14 | trimethoprim resistant dihydrofolate reductase dfr | antibiotic target replacement | diaminopyrimidine antibiotic |
| 8 | OXA-9 | OXA beta-lactamase | antibiotic inactivation | penam |
| 8 | aadA | ANT(3'') | antibiotic inactivation | aminoglycoside antibiotic |
| 8 | KPC-3 | KPC beta-lactamase | antibiotic inactivation | monobactam, carbapenem, cephalosporin, penam |
| 8 | AAC(6')-Ib' | AAC(6') | antibiotic inactivation | aminoglycoside antibiotic |

**Figure S2.**
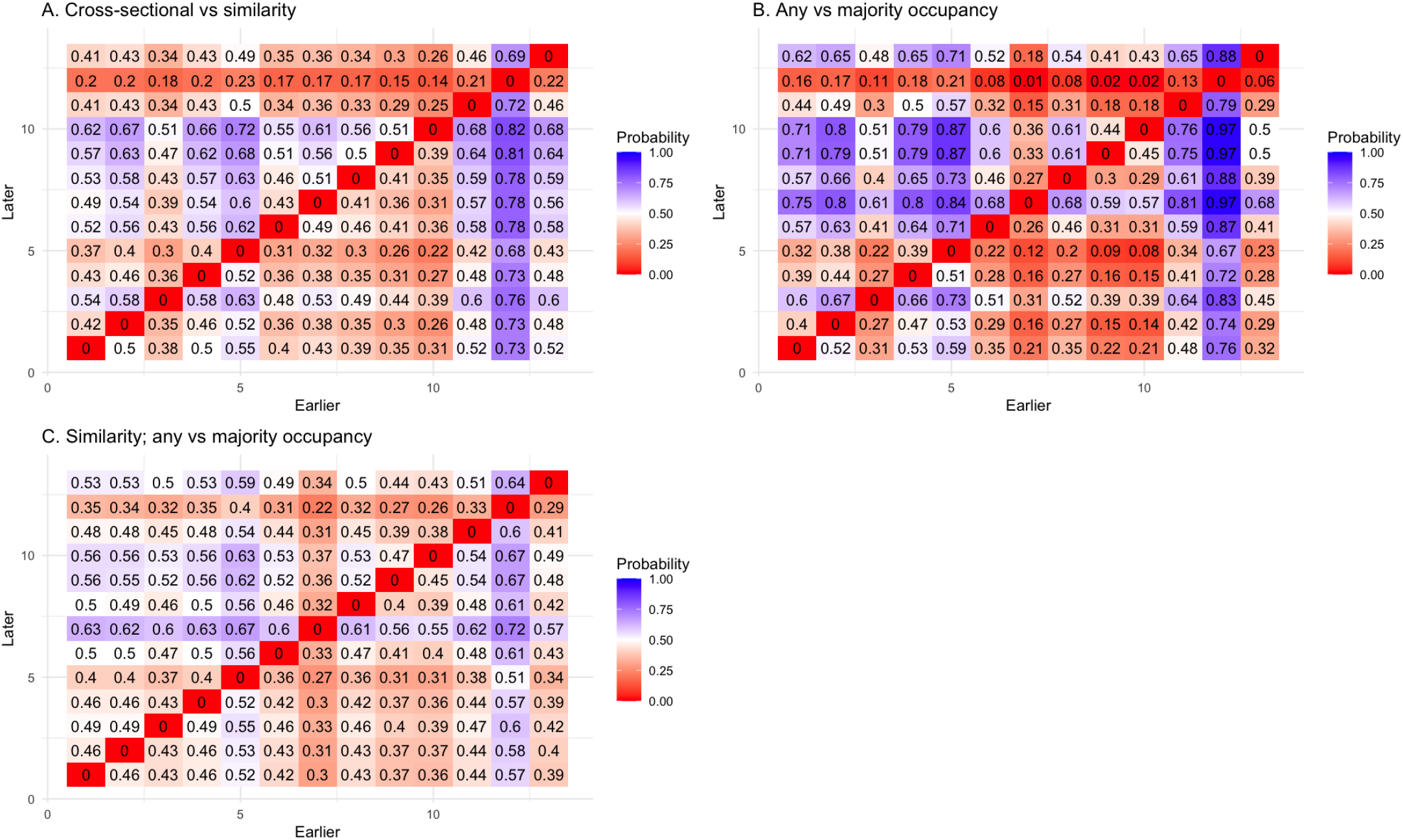
Severe malaria case study – comparing pairwise orderings. 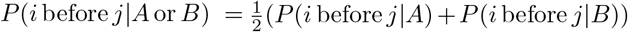 for ordering matrices *A* and *B* that come from different protocol choices. High-value (blue) elements denote a high probability that feature *i* (horizontal axis) is acquired before feature *j* (vertical axis), across the different protocol choices given. Low-value (red) elements denote a low probability (i.e. *j* is likely acquired before *i*).

**Figure S3.**
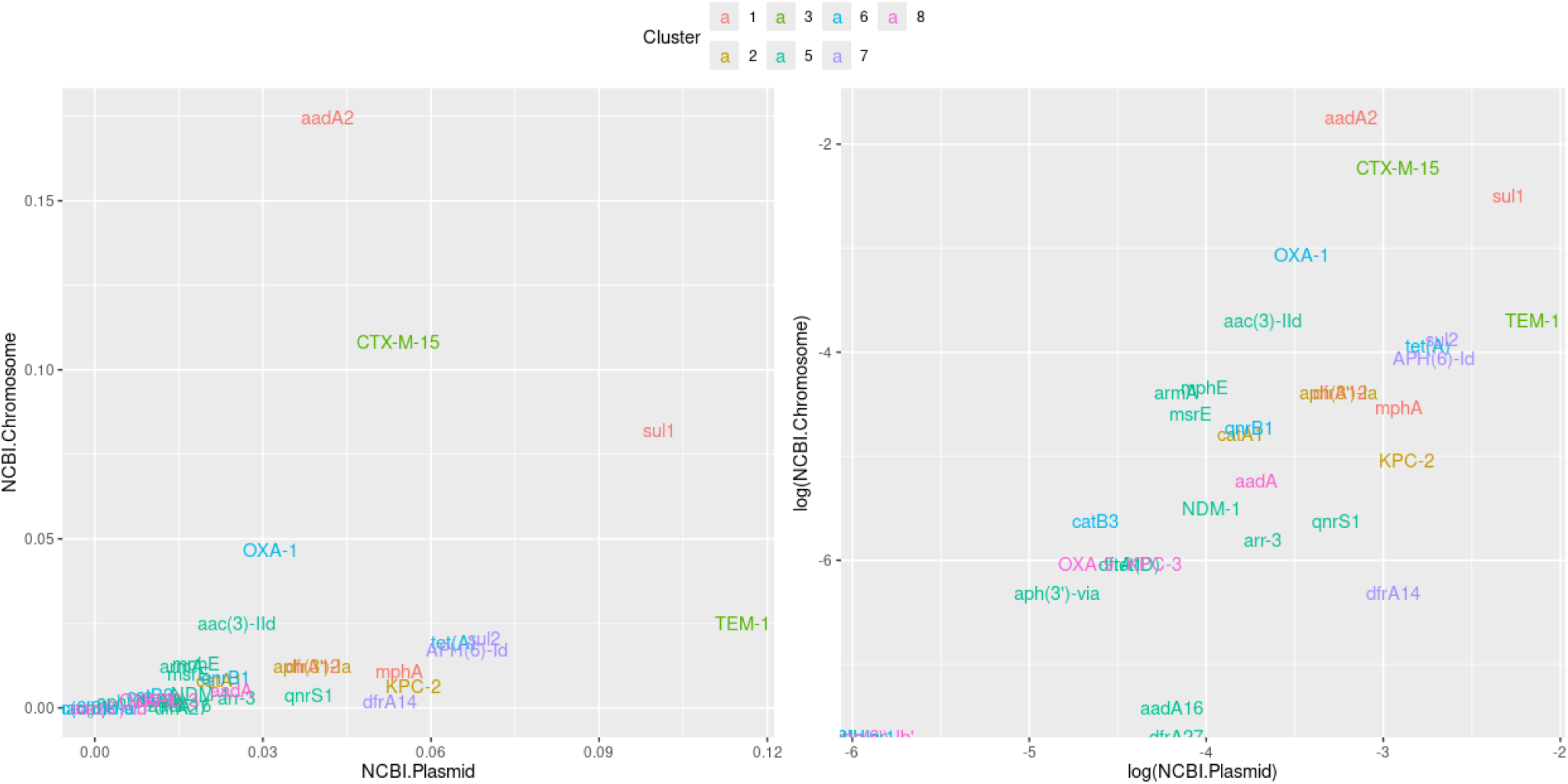
Location of genetic features in Klebsiella AMR case study. The percentage prevalence of the genetic features in Klebsiella pneumoniae isolates in either the NCBI plasmid or NCBI chromosome database. Mapped by CARD with the Resistance Gene Identifier [Alcock et al., 2023].

**Figure S4.**
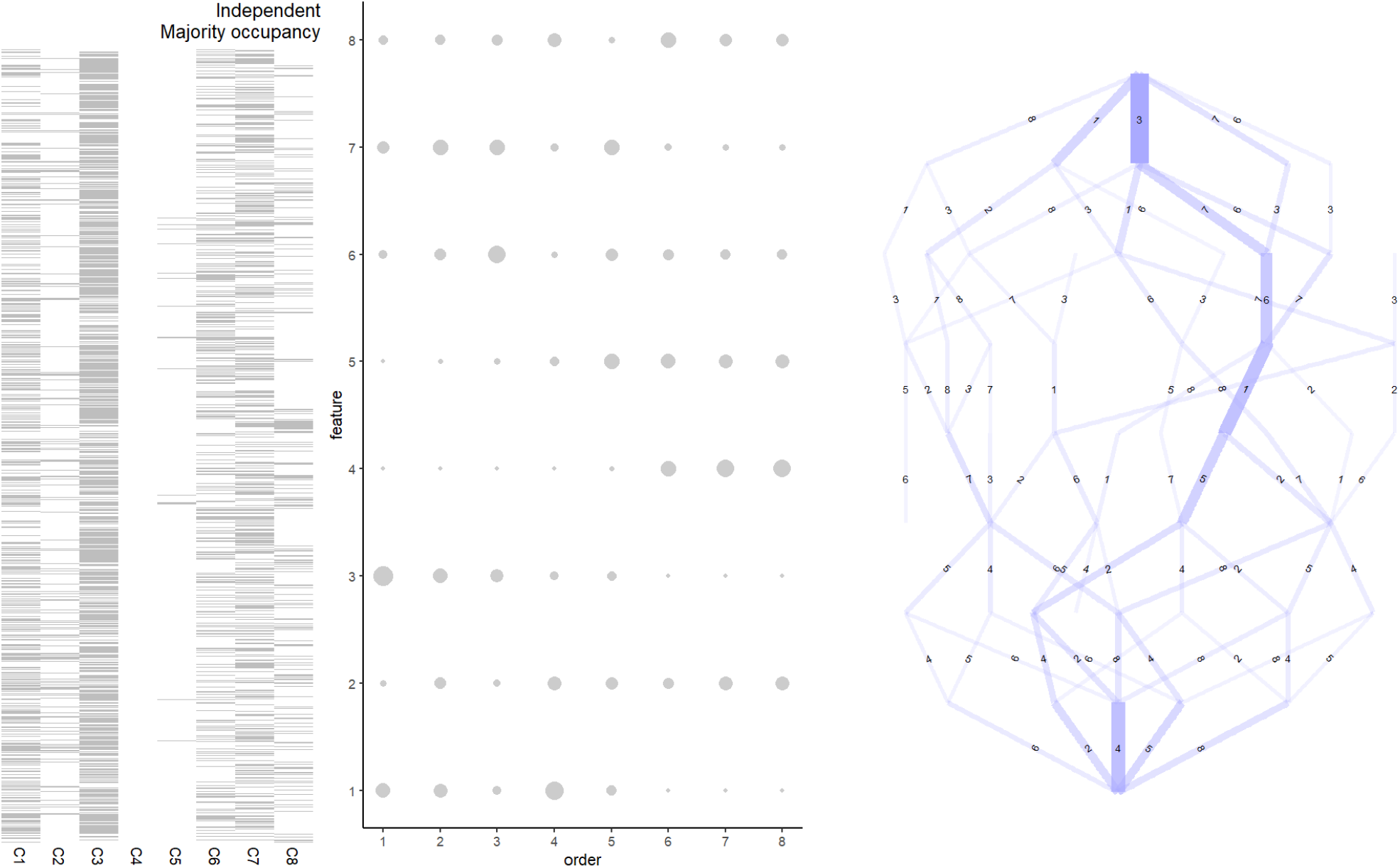
Klebsiella AMR case study with ‘majority’ occupancy rule.

**Figure S5.**
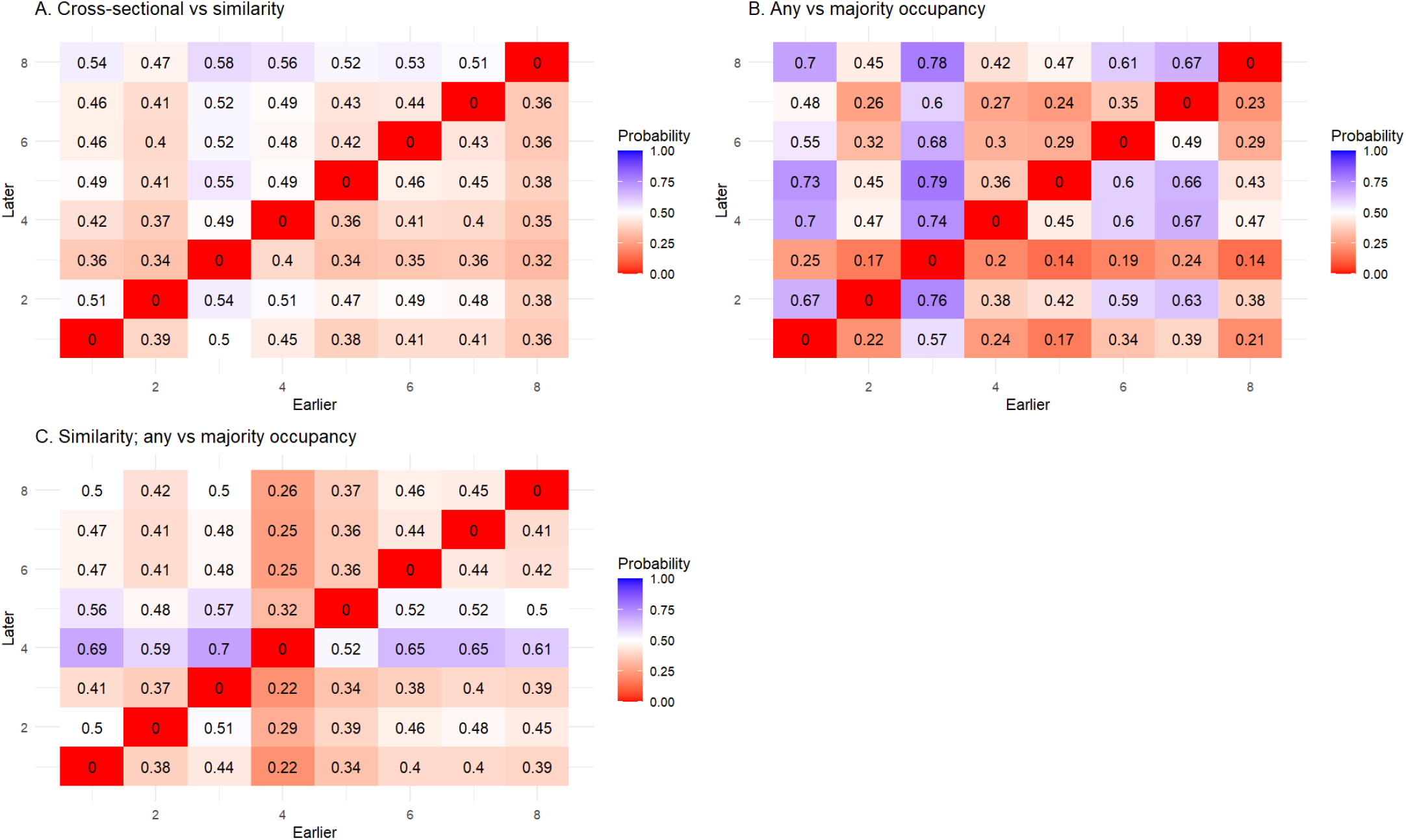
Klebsiella AMR case study – comparing pairwise orderings. 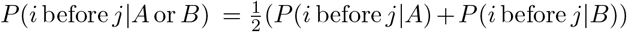 for ordering matrices *A* and *B* that come from different protocol choices. High-value (blue) elements denote a high probability that feature *i* (horizontal axis) is acquired before feature *j* (vertical axis), across the different protocol choices given. Low-value (red) elements denote a low probability (i.e. *j* is likely acquired before *i*).

**Figure S6.**
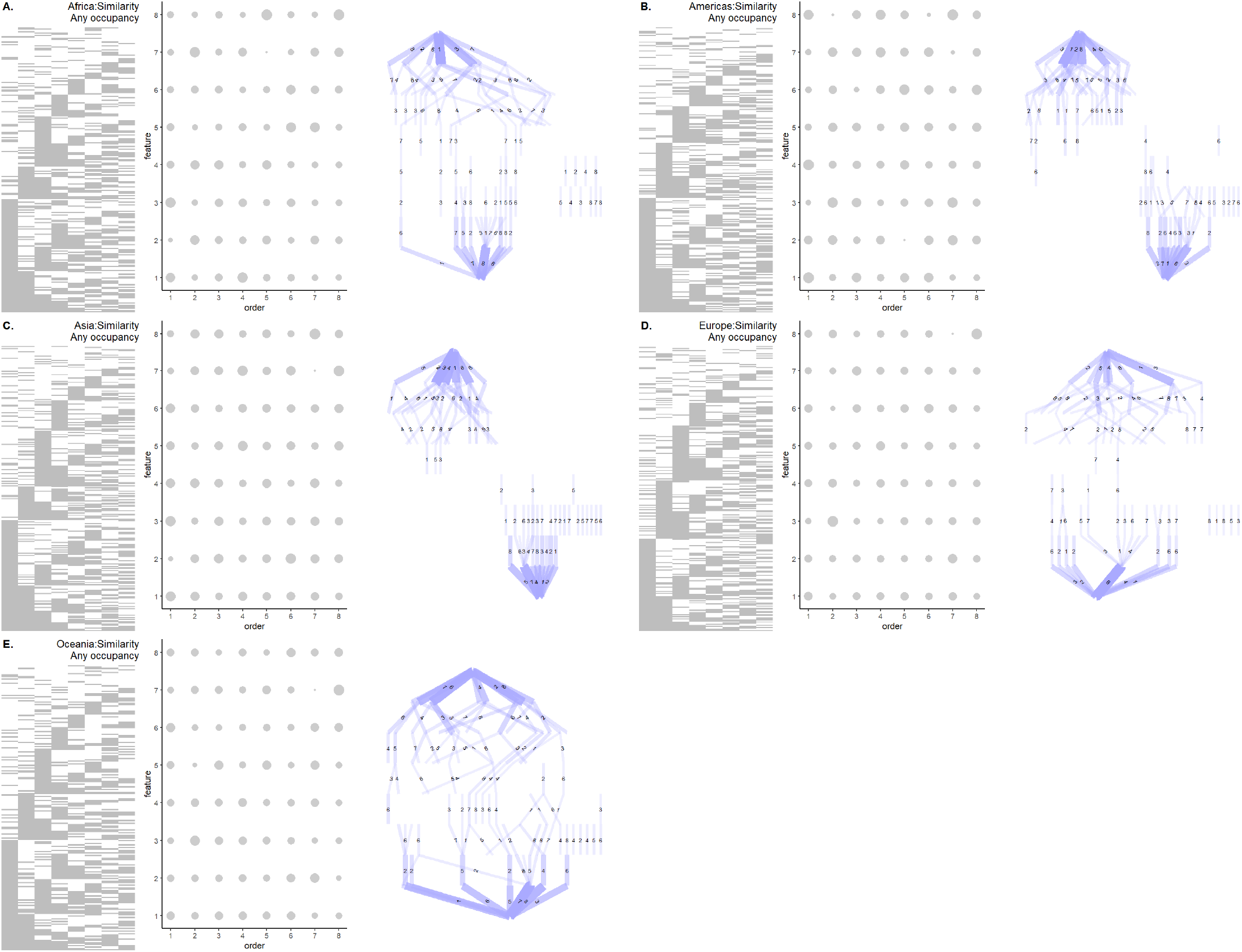
Klebsiella AMR case study across continents. Clustered data, ordering plots, and thresholded transition networks for subsets of the *Klebsiella* dataset by continent.

